# Proteome and phosphoproteome changes associated with prognosis in acute myeloid leukemia

**DOI:** 10.1101/796011

**Authors:** Elise Aasebø, Frode S. Berven, Sushma Bartaula-Brevik, Tomasz Stokowy, Randi Hovland, Marc Vaudel, Stein Ove Døskeland, Emmet McCormack, Tanveer S. Batth, Jesper V. Olsen, Øystein Bruserud, Frode Selheim, Maria Hernandez-Valladares

**Affiliations:** Department of Clinical Science, University of Bergen, Bergen, Norway; The Proteomics Facility of the University of Bergen (PROBE), University of Bergen, Bergen, Norway; The Department of Biomedicine, University of Bergen, Bergen, Norway; Department for Medical Genetics, Haukeland University Hospital, Bergen, Norway; Department of Biological Sciences, University of Bergen, Bergen, Norway; Centre for Cancer Biomarkers, Department of Clinical Science, University of Bergen, Bergen, Norway; Novo Nordisk Foundation Center for Protein Research, University of Copenhagen, Denmark

## Abstract

Acute myeloid leukemia (AML) is a hematological cancer that mainly affects the elderly. Although complete remission (CR) is achieved for majority of the patients after induction and consolidation therapies, nearly two-thirds relapse within a short interval. Understanding biological factors that determine relapse has therefore become of major clinical interest in AML. We utilized liquid chromatography tandem mass spectrometry (LC-MS/MS) to identify protein changes and protein phosphorylation events associated with AML relapse in primary cells from 41 AML patients at time of diagnosis. Patients were defined as relapse-free if they had not relapsed within a 5-year clinical follow-up after AML diagnosis. Relapse was associated with increased expression of RNA processing proteins and decreased expression of V-ATPase proteins. We also observed an increase in phosphorylation events catalyzed by cyclin-dependent kinases (CDKs) and casein kinase 2 (CSK2). The biological relevance of the proteome findings was supported by cell proliferation assays using inhibitors of V-ATPase (bafilomycin), CSK2 (CX-4945), CDK4/6 (abemaciclib) and CDK2/7/9 (SNS-032). While bafilomycin preferentially inhibited the cells from relapse patients, the kinase inhibitors were less efficient in these cells. This suggests that therapy against the upregulated kinases also could target the factors inducing their upregulation rather than their activity. In conclusion, our study presents markers that could help predict AML relapse and direct therapeutic strategies.

## Introduction

Acute myeloid leukemia (AML) is an aggressive and heterogeneous cancer.^1,2^ Acute promyelocytic leukemia (APL)^3^, one of the variants of AML, has a favorable prognosis with up to 95% of patients achieving complete remission (CR), and about 75% are cured by the combination of all-trans retinoic acid (ATRA) treatment and chemotherapy.^4^ Conversely, only 40-50% of patients with the non-APL variants achieve long-term AML-free survival after intensive chemotherapy often combined with allogeneic stem cell transplantation.^5–7^ Although the CR rate is 60-85% for younger patients (18-60 years old) who can achieve the most intensive treatment^8^, many patients achieving CR after the induction therapy die from later chemoresistant relapse.^9,10–14^ Design of better treatments that avert AML relapse is therefore of great importance.

Genetic aberrations and mutations (e.g on *CEBPA*, *NPM1* and *TP53*) are implemented in the diagnostic subclassification of AML patients and in risk adapted treatment procedures.^15–20^ However, liquid chromatography tandem mass spectrometry (LC-MS/MS)-based proteomics or phosphoproteomics can also be used for subclassification of patients with non-APL variants of AML.^21–26^ In the present study we therefore compared the proteome and phosphoproteome profiles of primary AML cells collected at the time of diagnosis for patients who later either became long-term leukemia-free survivors or suffered from chemoresistant relapse. Based on the proteomics and phosphoproteomics analysis of these two groups, we found common denominators in diagnostic samples which could be utilized for future treatments for patients that would later relapse.

## Materials and methods

### AML patients and sample collection

Written informed consent was obtained from all patients in accordance with the Declaration of Helsinki and the use of human leukemia cells for the present study was approved by the Regional Ethics Committee (REK III Vest 2013-634). Primary AML cells were collected from the peripheral blood of 41 patients with relatively high levels of circulating leukemia blasts at the time of diagnosis (i.e. >15 × 10^9^ AML cells/L, >80% of circulating leukocytes). Highly enriched AML cell populations (>95%) could then be prepared by density gradient separation alone (Lymphoprep, Axis-Shield)^27^, and these enriched cells were then cryopreserved.^28^ According to the patients’ relapse status after 5-year clinical follow-up post diagnosis, we classified 15 patients as long-term survivors (i.e. REL_FREE patients) and 26 patients as chemoresistant relapse patients (i.e. RELAPSE patients) after achieving CR. See **Figure 1A-B**, **Table 1** and supplemental Table 1 for a detailed description of these two groups. Nine AML patients external to the previously described cohort were used for validation studies and are described in detail in supplemental Table 2. Karyotyping and analysis of mutations of 54 myeloid relevant genes were available for most patients (supplemental Methods).

**Figure 1.**
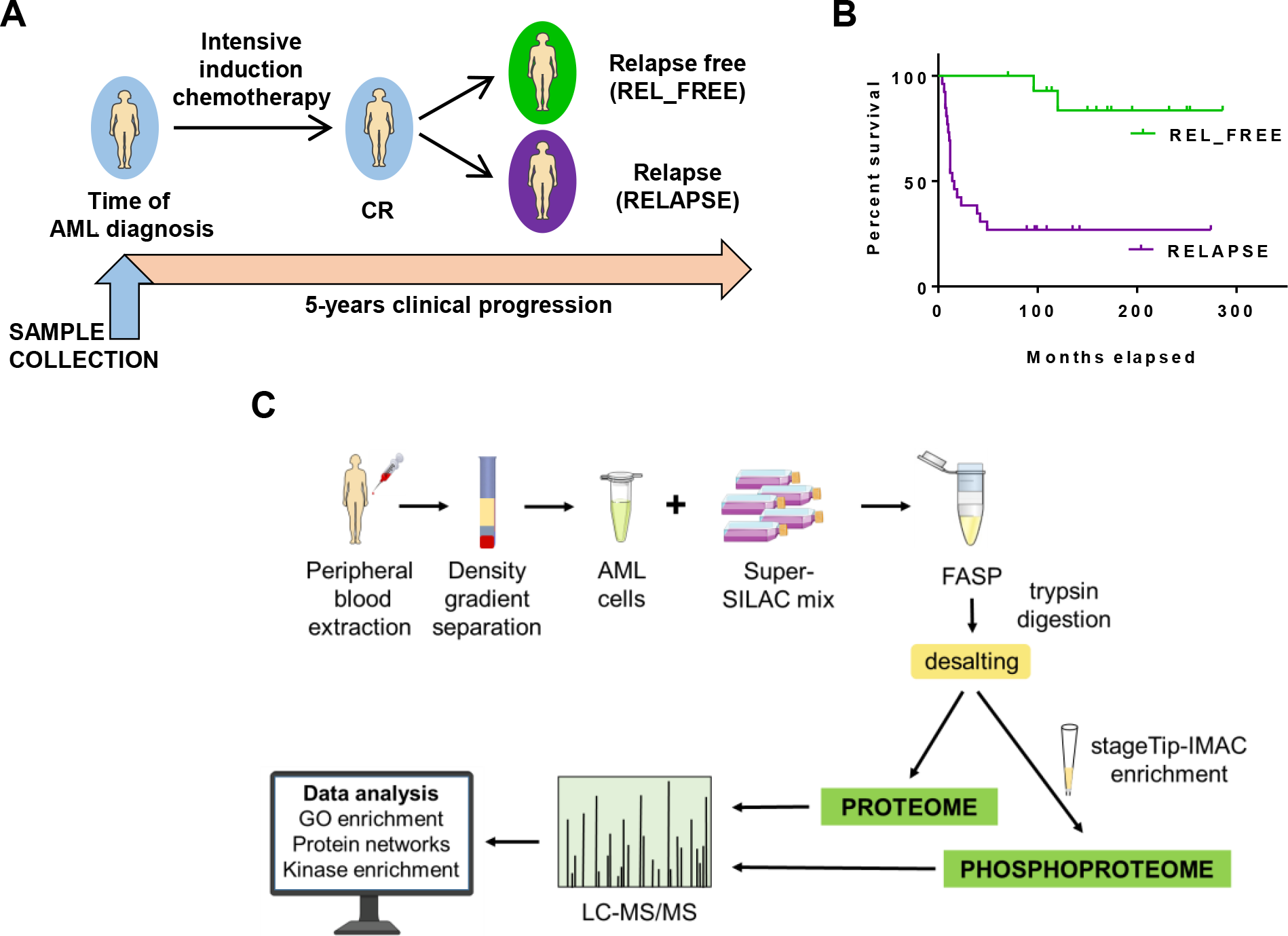
Overview of the RELAPSE and REL_FREE AML patient cohort, and the workflows for the proteome and phosphoproteome analysis of AML patient cells. **(A)** The study included AML cell samples from 26 RELAPSE and 15 REL_FREE patients collected at diagnosis time. All patients received intensive induction chemotherapy, consolidation therapy and achieved CR. **(B)** Survival plot for the patients included in each group. The observation time was at least 5 years. **(C)** AML sample preparation protocols for FASP-based proteome and phosphoproteome analysis followed by bio-computation and validation.

**Table 1.** Characteristics of the 41 AML patients at diagnosis

|  | REL_FREE | RELAPSE |
| --- | --- | --- |
| Age average (range) in years | 49.5 (36-65) | 50.5 (18-68) |
| Number of patients | 15 | 26 |
| **FAB classification** |  |  |
| M0-M1 | 1 | 11 |
| M2 | 0 | 2 |
| M4-M5 | 14 | 12 |
| uncertain | 0 | 1 |
| **FLT3** |  |  |
| WT | 14 | 14 |
| ITD | 1 | 8 |
| ND | 0 | 4 |
| **NPM1** |  |  |
| WT | 6 | 16 |
| Ins | 8 | 7 |
| ND | 1 | 3 |

FAB: French-American-British
ND indicates not determined; and WT, wild type
ITD: internal tandem duplication
Ins: a 4 bp-insertion/duplication

### AML super-SILAC mix

An AML spike-in reference was generated by combining Arg6- and Lys8-labeled protein cell lysates from five heterogeneous AML-derived cell lines, according to the super-SILAC (Stable Isotope Labeling with Amino acids in Cell culture) mix approach.^29,30^ The spike-in reference was added to each patient sample in an 1:2 ratio (w:w; super-SILAC mix:AML patient sample) for SILAC-based quantitation.

### Patient sample preparation for proteomic and phosphoproteomic analysis

Sample preparation of patient cell lysate in 4% sodium dodecyl sulfate (SDS)/0.1 M Tris-HCl (pH 7.6) and immobilized metal affinity chromatography (IMAC) has been described earlier.^31^ Briefly, 20 μg of each patient lysate was prepared both as 1) a label-free sample and 2) mixed with 10 μg of the super-SILAC mix for proteomic analyses, and processed according to the filter-aided sample preparation (FASP) protocol^31,32^ (supplemental Methods; **Figure 1C**). The super-SILAC spiked peptide samples were fractionated using styrenedivinylbenzene-reversed phase sulfonate (SDB-RPS) plugs (Empore, 3M).^33^ The phosphoproteomics samples (64-1121 μg range) were mixed with the super-SILAC mix at the ratio described before, FASP processed and enriched for phosphopeptides using the IMAC procedure.

### Nanoflow LC-MS/MS

Peptide sample preparation prior to LC-MS/MS and settings of the LC-MS/MS runs on a Q Exactive HF Orbitrap mass spectrometer coupled to an Ultimate 3000 Rapid Separation LC system (Thermo Scientific) were conducted as described earlier for global proteomics samples^34^ and phosphoproteomics^31^, and detailed in the supplemental Methods.

### Data and bioinformatics analysis

LC-MS/MS raw files from RELAPSE and REL_FREE samples were processed with MaxQuant software version 1.5.2.8^35,36^ (supplemental Methods). The spectra were searched against the concatenated forward and reversed-decoy Swiss-Prot Homo sapiens database version 2018_02 using the Andromeda search engine.^37^ The LC-MS/MS raw files and MaxQuant output files have been deposited to the ProteomeXchange consortium via the PRIDE partner repository ^38,39^ with dataset identifier PXD014997 (Reviewer account details, username: and password: jjAkVBC6). The Perseus 1.6.1.1 platform was used to analyze and visualize protein groups and phosphosites.^40^ MaxQuant-normalized SILAC ratios were inverted, log_2_ transformed and normalized again using width adjustment. Proteins and phosphosites (localization probability > 0.75) with at least five SILAC ratios in each patient group were selected for two-sample unequal variance *t*-test and *Z*-statistics^41^ to find significantly different fold change (FC) for proteins and phosphosites between the RELAPSE and REL_FREE groups.

Hierarchical clustering of significantly differential proteins and phosphosites was done with Perseus using the Pearson correlation function and complete linkage. Gene ontology (GO) analysis of biological process, molecular function and cell compartment terms was performed using a GO tool.^42^ The most significantly over-represented GO terms with false discovery rate (FDR) < .05 were displayed in bar plots in Prism8 (GraphPad). Venn diagrams were made with Venny 2.1 (*Oliveros, J.C.*, http://bioinfogp.cnb.csic.es/tools/venny/index.html). Gene Set Enrichment Analysis (GSEA) (http://software.broadinstitute.org/gsea/index.jsp) was performed against the Hallmark gene set collection of Molecular Signatures Database v6.2.^43^

Phosphosite and kinase prediction analyses were carried out as described in detail in supplemental Methods. Protein-protein interaction (PPI) networks were obtained by using the STRING database version 10.5 with interactions derived from experiments and databases at a high confidence score of 0.7.^44^ Networks were visualized using the Cytoscape platform version 3.3.0.^45^ The ClusterONE plugin was used to identify protein groups of high cohesiveness.^46^

### Enrichment analysis of transcription proteins binding sites

Putative binding sites of transcription proteins were found analyzing public available ChIP-seq data on K562 cell line at the ENCODE project.^47^ ENCODE accession IDs of files are provided in supplemental Table 3. Data analysis workflow was performed as detailed in supplemental Methods.

### Cell proliferation assay

The cytokine-dependent *in vitro* proliferation of enriched AML cells was analyzed by a 3H-thymidine incorporation assay (see supplemental Methods).^48^ The median of triplicate determinations was used in all analyses. Mann-Whitney U test (SPSS Statistics version 25, IBM) was employed for analysis of differences between different groups and Wilcoxon’s matched-pair signed rank test (SPSS) for analysis of differences between paired samples.

### Western blots

Western blots for three RELAPSE and three REL_FREE patient cells used in the cell proliferation assay and incubated with kinase inhibitors at 37 °C for 15 minutes were performed as described in supplemental Methods.

## Results

### The protein abundances of rRNA processing proteins and V-ATPase subunits differ between RELAPSE and REL_FREE patients

We compared the proteome profiles of AML cells derived from 26 RELAPSE and 15 REL_FREE patients at the time of diagnosis. We quantified 6781 proteins, of which 5309 had a quantitative value in at least five patients in each group. The *t*-test based statistical analysis resulted in 351 differentially expressed proteins, 210 proteins were upregulated and 141 were downregulated in the RELAPSE relative to REL_FREE group (supplemental File 1). The proteome of patient-derived cell lysates were also processed and quantified using label-free quantification (LFQ) for validating the super-SILAC based proteome quantitation (supplemental Data, supplemental Figure 1, 2 and 3 and supplemental File 1). The RELAPSE vs REL_FREE FCs of the 4041 proteins quantified in both experiments had good correlation (Pearson *R* = .72; supplemental Figure 2).

Hierarchical clustering of the 351 differential proteins grouped the patients into two main clusters (**Figure 2A**), which corresponded to the RELAPSE and REL_FREE samples, although a distinct separation was not obtained. GO enrichment analysis showed that ATPase activity (coupled to transmembrane movement of ions, rotational mechanism), proton-exporting ATPase activity, phagosome acidification and iron ion transport (**Figure 2B**) were more abundant processes in the REL_FREE patients. These terms include proteins of the vacuolar (V)-H^+^-ATPase, an ATP-dependent proton pump in cellular membranes such as lysosomes and endosomes. The proteins enriched in this group were primarily located in the cytosol or extracellular organelles such as vesicles. In contrast, proteins involved in ribosomal processes such as rRNA metabolic process, ncRNA processing and ribonucleoprotein complex biogenesis were clearly higher in the RELAPSE patients, where 46% of the increased proteins were annotated to the nucleolus in the cellular compartment analysis (**Figure 2B**).

**Figure 2.**
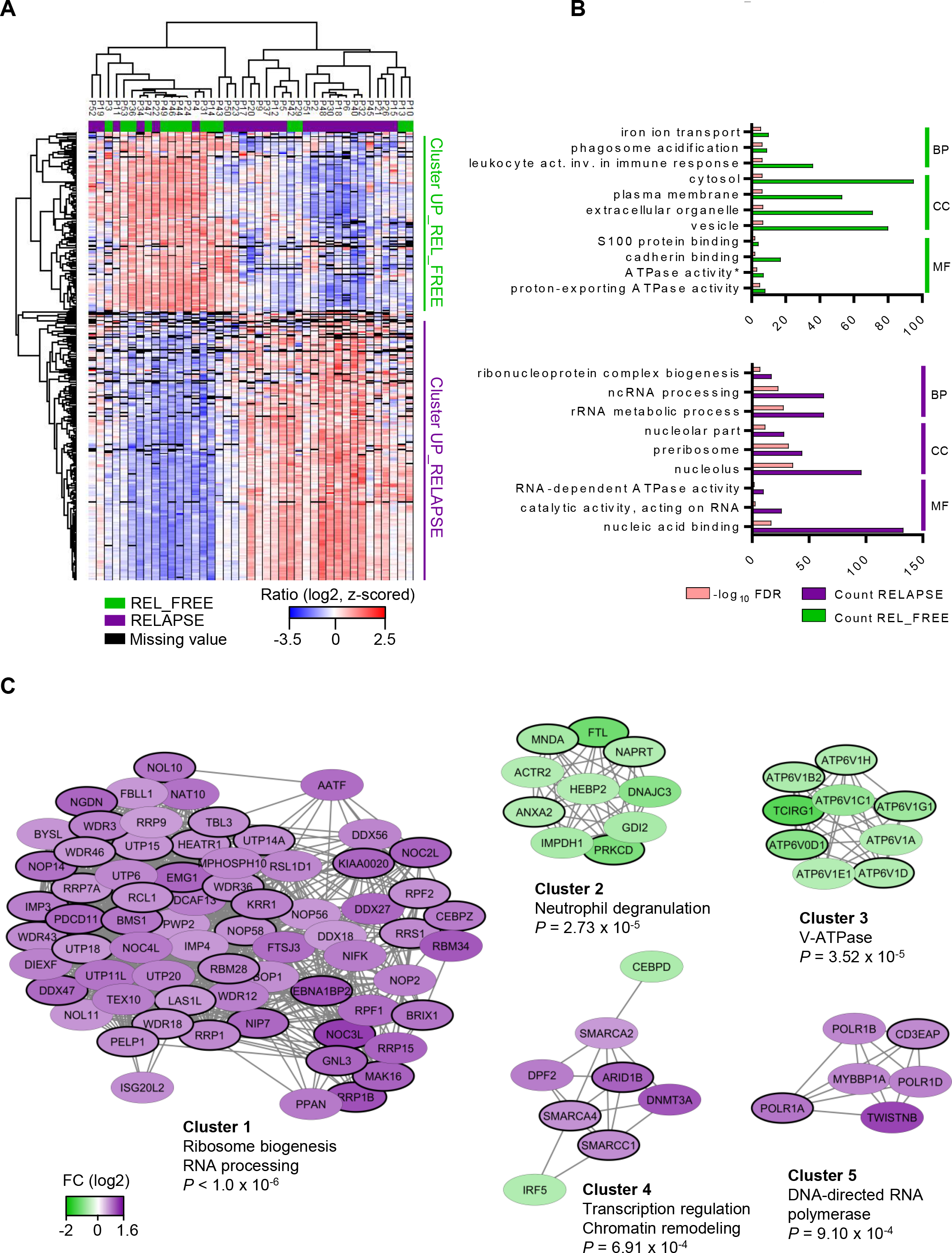
The global AML cell proteome show increased abundance of rRNA processing proteins and decreased abundance of V-ATPase subunits for patients who relapse. **(A)** Hierarchical clustering of the patients (P1-P52) based on protein expression (SILAC log_2_ ratio) of the 351 proteins with significantly different regulation in AML cells from REL_FREE (green) and RELAPSE (purple) patients. Two vertical main clusters were observed, one dominated by proteins with higher abundance in mostly REL_FREE patients (Cluster UP_REL_FREE) and the other by proteins with higher abundance in RELAPSE patients (Cluster UP_RELAPSE). **(B)** Gene Ontology (GO) analyses of the two protein clusters were performed to reveal enriched biological processes (BP), cellular compartments (CC) and molecular functions (MF) in the two protein clusters. The number of proteins associated to a specific GO term (count) and −log_10_ FDR of the top significant GO terms are shown. *Full GO term: ATPase activity, coupled to transmembrane movement of ions, rotational mechanism. **(C)** PPI networks of the 351 proteins from the STRING database, visualized and analyzed with Cytoscape and ClusterONE, respectively. The five clusters with highest significance of cohesiveness are shown with the *P* value of a one-sided Mann-Whitney U test. The protein nodes are colored according to their RELAPSE/REL_FREE log_2_ FC, i.e. purple indicates increased abundance in the RELAPSE group and green increased abundance in the REL_FREE group.

Moreover, a PPI network of the differentially expressed proteins (**Figure 2C**) confirmed the increased abundance of proteins involved in rRNA processing and ribosome biogenesis in relapsed patients (cluster 1; 69 proteins). RRP1B (*P* < .0002, FC 1.09) and AATF (*P* < .0002, FC 0.91) were the most significant proteins in this cluster. AATF (also known as CHE1) is participating in several cellular pathways, such as 40S ribosomal subunit synthesis in complex with NGDN (*P* = .0014, FC 0.95) and NOL10 (*P* = .0007, FC 0.85)^49^, and high AATF expression has been linked to variants of leukemia^50,51^. Other proteins in this cluster were CEBPZ, a transcription factor in the CEBP family, several WD repeat-containing proteins (e.g. WDR3, WDR18, WDR36), probable ATP-dependent RNA helicases (e.g. DDX18, DDX27, DDX56) and members of the C/D box small nucleolar ribonucleoprotein (snoRNP) complex (NOP56 and NOP58) which guides 2’O methylation of the rRNA. Proteins involved in neutrophil degranulation dominated cluster 2 and nine subunits belonging to the V-ATPase complex were identified in cluster 3 (**Figure 2C**), all with higher abundance in the REL_FREE patients. Regulated V-ATPase subunits (cluster 3) and proteins participating in rRNA processing and ribosomal biogenesis (cluster 1) are detailed in supplemental File 2. Cluster 4 and 5 contained proteins involved in chromatin remodeling and RNA polymerase I subunits respectively, which were upregulated in the RELAPSE patients.

The GSEA analysis against the Hallmark gene set collection, conducted to identify classes of genes overrepresented in the complete dataset, resulted in HALLMARK_MYC_TARGETS_V2 (systematic name: M5928) as the only significant gene set (FDR q-value = .0461) (supplemental Figure 4). This gene set was upregulated in the RELAPSE group and consisted of 58 genes.^52^ A total of 15 out of the 30 proteins who contributed most to the gene set enrichment results (i.e. the leading edge subset) were found in cluster 1 in **Figure 2C**.

### Differential CDK, CSK2 and PRKCA/D kinase activities between RELAPSE and REL_FREE patients

We constructed a dataset comprising 12309 identified and quantified class I protein phosphorylation sites from 3003 proteins of 26 RELAPSE and 15 REL_FREE patients. Serine phosphorylation made up majority of the identified phosphosites (89.7%) while phosphorylation on threonine and tyrosine (10.0 and 0.3%, respectively) comprised the rest. We identified 274 differentially regulated phosphorylated sites based on statistical analysis of 5634 phosphosites which were quantified in at least five patients in each group (supplemental File 3).

Hierarchical clustering using these 274 phosphosites divided the phosphoproteome of RELAPSE and REL_FREE patients (**Figure 3A**). Two clusters, one containing 138 phosphosites and another with 136 phosphosites, were upregulated and downregulated in the RELAPSE relative to REL_FREE patients, respectively.

**Figure 3.**
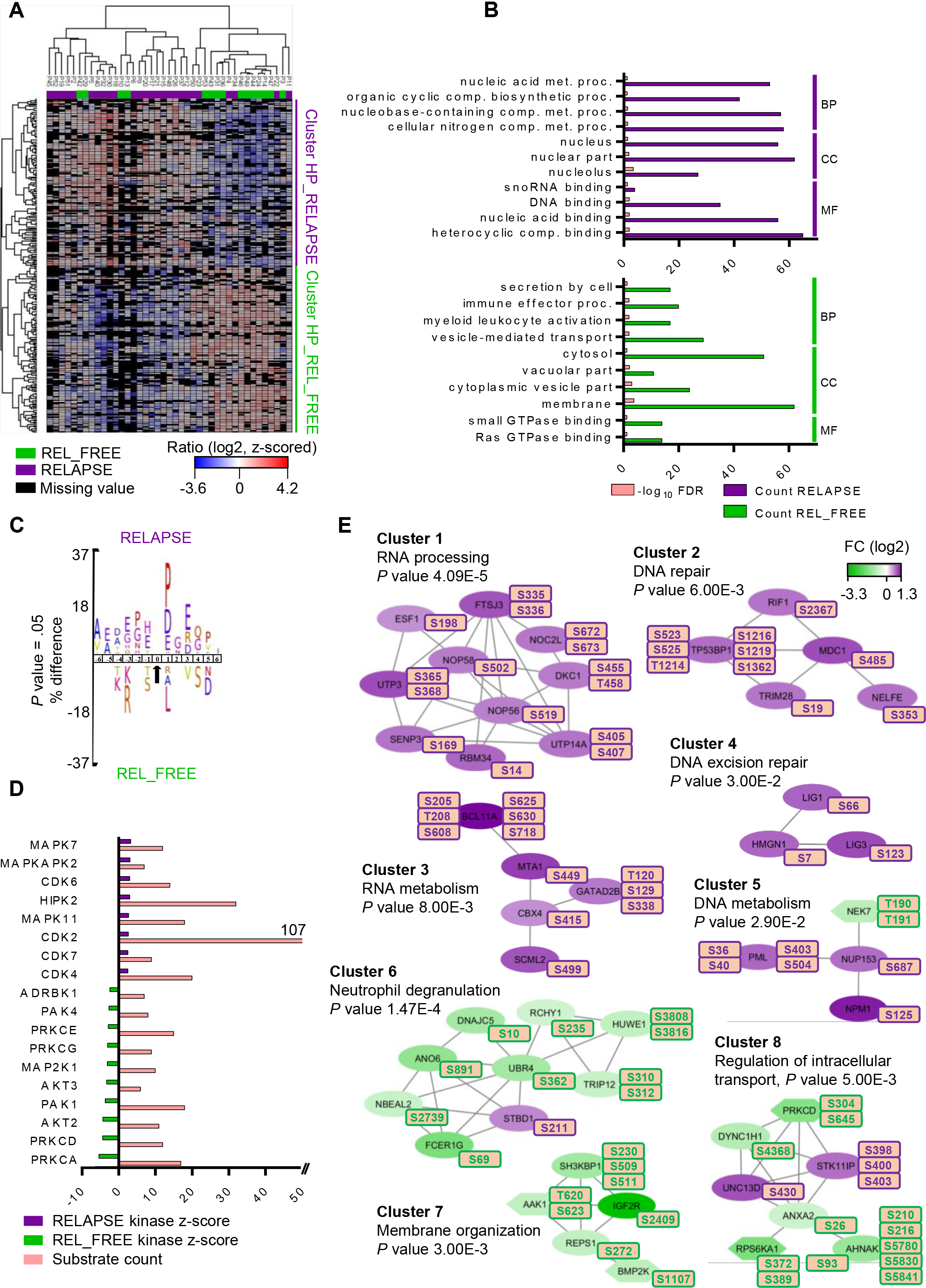
The AML RELAPSE phosphoproteome is enriched in CDKs substrates and RNA processing CSK2 targets. **(A)** Hierarchical clustering of 274 differentially regulated phosphorylation sites revealed two clusters: HP (High Phosphorylation)_RELAPSE (in purple) and HP_REL_FREE (in green). The SILAC log_2_ ratio scale and color code is also shown. **(B)** GO analyses for enriched BP, CC and MF was conducted for each protein cluster, and the count and −log_10_ FDR of the top significant GO terms are displayed. **(C)** Sequence motif analysis of the ± six amino acids flanking the differentially regulated phosphorylation sites for either cluster. **(D)** KSEA of differentially regulated and unregulated phosphorylation sites. The kinase z-score (X axis) is the normalized score for each kinase (Y axis), weighted by the number of identified substrates. **(E)** Networks of PPI based on STRING database and visualized in Cytoscape after ClusterONE analysis. Significance of networks of high cohesiveness is shown with the *P* value of a one-sided Mann-Whitney U test. The differentially regulated phosphorylation site(s) is shown next to each protein. FC of phosphorylation are color-coded; purple-colored proteins showed a higher phosphorylation in the RELAPSE group and green-colored proteins showed a higher phosphorylation in the REL_FREE group. Kinases are specifically distinguished using hexagon shapes.

Cellular component GO analysis revealed an enrichment of upregulated nuclear phosphoproteins for RELAPSE patients, whereas cytoplasm, cytosol, membranes and vacuolar structures were enriched in the REL_FREE group (**Figure 3B**, supplemental File 3). Immune effector process, myeloid leukocyte activation and GTPase binding were the biological process and molecular functional GO terms enriched in this group.

To identify protein kinases differentially activated in the two groups we performed phosphorylation site motif analysis (IceLogo).^53^ We found a strong bias towards acidic amino acids such as glutamic acid (E) in close proximity to the differentially phosphorylated sites in the RELAPSE group when compared to the unregulated phosphosites (supplemental Figure 5A), suggesting a higher CSK2 activity in this group. Using differentially phosphorylated sites in the REL_FREE group, the analysis identified a basophilic KXpS/pT motif, suggesting a moderate activation of PRKCA and PRKCD (supplemental Figure 5B). Moreover, proline-directed motifs for phosphorylation by MAPK and CDK kinases were underrepresented in the REL_FREE group. When we directly compared the sequences surrounding the differentially regulated phosphorylation sites in the RELAPSE and REL_FREE groups, higher CSK2, MAPK and CDK kinase activation was observed in the RELAPSE, and higher PRKCA/D in the REL_FREE group, confirming the motif analysis (**Figure 3C**, supplemental File 4). Furthermore, the KSEA analysis^54,55^ (which is based on phosphorylation FCs to estimate kinase’s activity) confirmed the higher activity of several MAPK kinases and CDK kinases in the RELAPSE group and the higher activity of PRKCA, PRKCD and AKT kinases in the REL_FREE group (**Figure 3D**). We found 9 phosphosites on 6 different protein kinases in the data set of 274 differentially regulated phosphorylation sites. Seven of the nine phosphosites (including those on PRKCD, PAK2, MAP3K3, BMP2K and NEK7 kinases) were found to be upregulated in the REL_FREE group relative to the RELAPSE group. In particular, sites on the kinase domain activation loop of NEK7 (T190 and T191) were more abundant in the REL_FREE group. Conversely, phosphorylation of FLT3 on S759 and S762 was higher in the RELAPSE group relative to the REL_FREE group.

Several PPI networks of significant cohesiveness were found after ClusterONE analysis based on STRING interactions of differentially phosphorylated proteins (**Figure 3E**). The most significant network (cluster 1) consisted of ten phosphoproteins with higher phosphorylation in RELAPSE patients, most of them being RNA binding proteins. A sequence logo analysis^56^ of the amino acids surrounding the 15 phosphosites (corresponding to the ten proteins) suggested a higher activation of CSK2 in this cluster (supplemental Figure 6A). Other significant clusters included DNA binding proteins (cluster 2 and 4) and proteins involved in nucleic acid metabolism (cluster 3 and 5). A sequence logo analysis of the amino acids surrounding the phosphosites in clusters 2, 3 and 5 showed an enrichment of MAPK substrates (supplemental Figure 6B-D), although NPM1 S125 in cluster 5 is phosphorylated by CSK2^57^, and amino acid sequences around HMGN1, LIG1 and LIG3 phosphosites in cluster 4 suggested the involvement of other kinases such as CSK2 and CDK^58,59^. Finally, transcription factor BCL11A, which functions as a myeloid proto-oncogene, and DNA repair protein TP53BP1 showed higher phosphorylation on multiple sites in the RELAPSE group relative to the REL_FREE group.

Conversely, PPI networks that showed higher phosphorylation in the REL_FREE group were involved in neutrophil degranulation (cluster 6), membrane organization (cluster 7) and intracellular transport (cluster 8). Sequence logo analysis of the phosphosite motifs in these clusters suggested a higher activation of PRKCD/A kinases (supplemental Figure 6E-G).

To further elucidate the molecular source of AML relapse in these patients, we utilized the proteome and phosphoproteome data in order to determine transcription factors responsible for driving relapse. Specifically, proteins involved in DNA transcription that were regulated at both the proteome or phosphoproteome level in the RELAPSE group were compared to ChIP-seq data from the K562 cell line (stored in the ENCODE database) in order to find potential binding sites of transcription factors (supplemental Table 3). We identified six transcription factors (AFF1, CEBPZ, ETV6, IKZF1, PML and TRIM28) that showed predicted regulation of proteins involved in DNA and RNA binding proteins, damaged DNA binding proteins, ribosome and carcinogenesis pathways based on GO molecular function terms and KEGG pathways analysis (supplemental Figure 7).

To explain the generally increased phosphosite regulation in the RELAPSE group, we compared the protein expression to the 274 differentially regulated phosphosites (corresponding to 169 phosphoproteins). We found that 107 (63%) phosphoproteins were not significantly changed at the protein level (supplemental File 3). Among them, we noticed all the phosphoproteins of the DNA repair network (cluster 2) in **Figure 3E** (supplemental Figure 8). In contrast, we spotted 34 proteins significantly regulated at both protein and phosphosite level, including 6 phosphoproteins of the RNA processing network (cluster 1) in **Figure 3E** (supplemental Table 4 and supplemental Figure 9). For majority of these phosphosites (including FLT3 and PML), the phosphorylation levels correlated closely with their protein expression levels. Taking together, while RELAPSE AML cells had higher expression of several phosphorylation-prone proteins, nearly 50% of the differentially regulated phosphorylation sites could not be explained by protein expression changes, suggesting increased kinase-specific phosphorylation.

### V-ATPase, CSK2, CDK2/7/9 and CDK4/6 inhibitors affect the proliferation of AML cells

To explore the potential of V-ATPase inhibition in AML, we tested the effect of the V-ATPase inhibitor bafilomycin A1 (BafA1) on the growth of AML cells from four REL_FREE and seven RELAPSE patients with high or low abundance of V-ATPase subunits, respectively (supplemental Figure 10A-C). The proliferation was significantly higher in the untreated control RELAPSE cells compared to REL_FREE cells (supplemental Figure 10D). BafA1 at 10 nM decreased significantly (*P* = .018) the proliferation of RELAPSE cells, whereas the proliferation of the REL_FREE cells was not significantly altered (**Figure 4A**).

**Figure 4.**
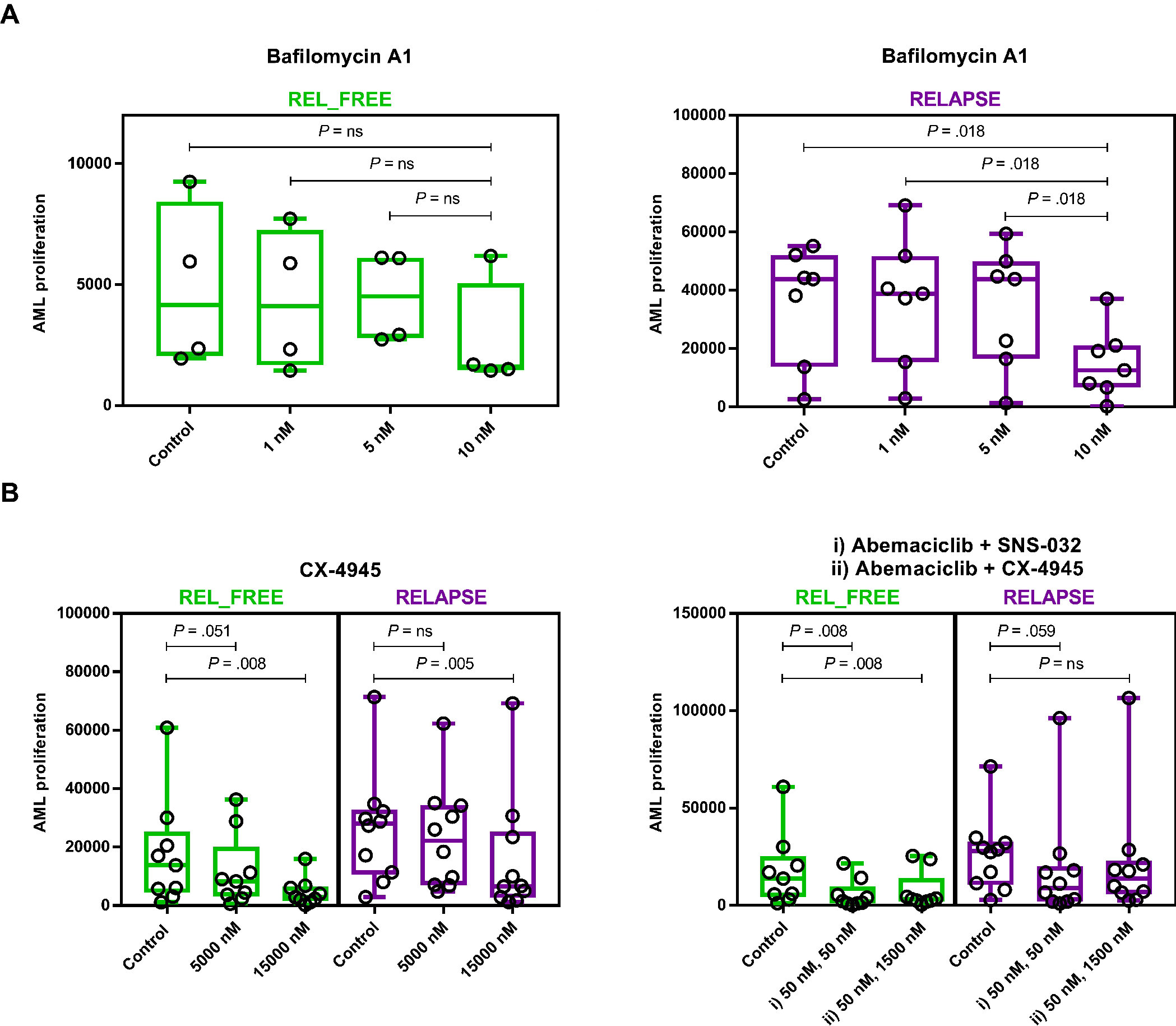
The effect of inhibitors of V-ATPases, CSK2 or CDKs on AML cell proliferation. AML patient cells were treated for six days with the indicated inhibitors. The thymidine incorporation-based proliferation measurements are presented as dots, while the median proliferation (in counts per minute) and min to max are presented as box plots. Significance of inhibitor treatment versus untreated control was found using the Wilcoxon matched-pair signed rank test. Non-significant results are indicated with “*P* = ns”. **(A)** Cells from four REL_FREE and seven RELAPSE patients treated with V-ATPase inhibitor bafilomycin A1 (BafA1). The RELAPSE, but not the REL-FREE, patients had decreased proliferation (measured after 6-7 days of culture) when treated with 10 nM BafA1 inhibitor compared to control, 1 and 5 nM. No significant change was observed between control and 1 nM or 5 nM BafA1 inhibitor for neither groups **(B)** Cells from nine REL_FREE and ten RELAPSE patients (five and four of them were external to the patient cohort, respectively) were treated with CSK2, CDK2/7/9 and CDK4/6 kinase inhibitors in cell proliferation assays, alone or in combination. The CSK2 inhibitor CX-4945 was given in 0 (i.e. control), 5000 and 15000 nM alone (left plot). The CDK4/6 inhibitor Abemaciclib (50 nM) was given in combination with i) the CDK2/7/9 inhibitor SNS-032 (50 nM) and ii) CX-4945 (1500 nM) (right plot).

In order to further investigate whether the activity of the predicted RELAPSE-activated kinases was essential for cell viability, we tested proliferation in AML cells from ten RELAPSE and nine REL_FREE patients in the presence of inhibitors of CSK2, CDK2/7/9, CDK4/6 and ERK1/2, alone or in combination (supplemental Figure 11A). As shown in **Figure 4B**, the CSK2 inhibitor CX-4945 at 5000 nM was a slightly more potent inhibitor of REL_FREE cells compared to RELAPSE cells. Similarly, the combination of CDK4/6 inhibitor abemaciclib (50 nM) with CDK2/7/9 and CSK2 inhibitors (SNS-032 50 nM; CX-4945 1500 nM) decreased cell proliferation significantly only for the REL_FREE cells (*P* = .008). The RELAPSE cells showed significant anti-proliferative effect only when exposed to a high concentration (15000 nM) of CX-4945.

Western blot analyses using lysates obtained from AML cells from three RELAPSE and three REL_FREE patients incubated with CDK2/7/9 or ERK1/2 inhibitors showed both higher CDK2 expression and higher phosphorylation of CDK2 on T160 in the RELAPSE group. The phosphorylation of ERK 1/2 was low and similar in the two patient groups (supplemental Figure 11B). These findings correlate with the phosphoproteome analyses showing higher relative abundance of phosphorylated CDK2 substrates, but not of ERK1/2 substrates, in the RELAPSE group.

## Discussion

Despite the patient heterogeneity, significant overall differences were observed at the proteome and phosphoproteome level for several biological processes between AML cells from RELAPSE and REL_FREE patients, harvested at the time of diagnosis. In particular, proteins involved in ribosome biogenesis and rRNA processing/regulation were more abundant and more strongly phosphorylated in RELAPSE patient cells. This finding was supported by the ChIP-seq data analysis from the K562 cell line where we found several transcriptions factors promoting transcription of genes for RNA polymerase (CEBPZ) and ribosome/ribosomal biogenesis (PML, AFF1, ETV6 and TRIM28). Deregulated RNA binding proteins and ribosome biogenesis can be drivers of cancer pathogenesis.^60,61^ Furthermore, the GSEA analysis revealed enrichment of MYC targets in RELAPSE patients. MYC is frequently overexpressed in AML as well as other leukemic malignancies, and seems to be a crucial transcriptional factor during hematopoiesis.^62,63^ MYC is also a regulator of ribosome biogenesis^64^, activates RNA pol I for rDNA transcription^65^ and induces snoRNA expression^66^. As MYC targets are enriched during relapse, our findings suggest that MYC, together with proteins involved in rRNA processing and ribosome biogenesis, are prognostic biomarkers for relapse and potential therapeutic targets in AML. Several drugs targeting rRNA synthesis, transcription or processing in hematological cancers have already been described.^67,68^

We observed decreased abundance of V-ATPase subunits in RELAPSE patients. The V-ATPase complex can modulate several intracellular signaling pathways with importance in AML (such as PI3K-mTOR^69,70^, NOTCH1^71^ and WNT^72^) by controlling acidification of intracellular compartments. When testing the effect of V-ATPase inhibitor on AML cells we observed that the RELAPSE derived cells with initially low abundance of V-ATPase subunits responded better to BafA1 V-ATPase inhibitor, thus, a potential relationship between low initial V-ATPase subunit abundance and chemoresistance needs further evaluation. V-ATPase is regarded as a potential therapeutic target in AML.^73^ Although material from only a limited number of patients was available for testing in the proliferation assay, differences in proteomic profiles were associated with differences in anti-proliferative pharmacological effects.

Alignment of sequences surrounding phosphorylation sites and KSEA analysis of our full phosphoproteomics dataset showed an enrichment of phosphorylated substrate groups of MAPK, CSK2, CDK, PAK, PRKC and AKT kinases. We additionally observed that RELAPSE cells had higher activity of CDK2 and higher expression based on immunoblotting assays compared to REL_FREE cells. As similarly demonstrated previously, KSEA of phosphoproteome data of AML-derived cells from 20 patients had similar enrichment of CSK2, CDK, PAK, AKT, ABL, SRC, MAPK, PRKA and PRKC kinases relative to normal peripheral blood stem CD34^+^ cells from 5 healthy donors.^55^ Thus, two independent phosphoproteomics studies on different AML patient cohorts have identified similar sets of kinase upregulations in AML.

The CSK2 inhibitor CX-4945 had a stronger anti-proliferative effect for REL_FREE cells than for RELAPSE cells when acting alone or in combination with CDK inhibitors. Further studies are required to determine whether RELAPSE cells owe their higher CSK2 inhibitor resistance to generally increased robustness or to lower turnover of CSK2-mediated phosphorylation. Comprehensive KSEA, sequence logos and antibody blotting analyses show that CSK2 and CDKs could be suggested as strong RELAPSE promoters. CSK2 in particular is a key regulator of several signaling networks and is overexpressed in many hematological cancers.^74^ The activity of CDK2 (and potentially also CDK4, CDK6, CDK7 and CDK9) in RELAPSE patients may further increase AML cell proliferation by triggering G1-S-phase transitions.^75^

Several phosphoproteins were multi-phosphorylated, like BCL11A with several phosphosites upregulated in the RELAPSE group (S205, T208, S608, S625, S630 and S718). BCL11A, a transcription factor associated with the BAF SWI/SNF chromatin remodeling complex, may play a role in leukemogenesis^76^ and its expression is associated with adverse outcome in AML.^77^ In a previous phosphoproteomics study with AML patients treated with the FLT3 inhibitor quizartinib, increased phosphorylation of BCL11A S630 was associated with nonresponsiveness.^24^ Taken together these results suggest that BCL11A phosphorylation contributes to chemoresistance in AML. TP53BP1, another multi-phosphorylated protein in RELAPSE patients, is a double-strand DNA break repair protein that is phosphorylated at multiple sites in response to DNA damage. Moreover, phosphorylation at S1219 by ATM kinase seems to be a response to ionizing radiation.^78^ The S523Q motif might also serve as substrate for ATM kinase and being part of the RIF1, a non-homologous end joining-mediated repair protein, recruitment process to DNA break sites.^79^

The stimulatory T190 and T191 phosphosites in the activation loop of NEK7 showed increased phosphorylation in the REL_FREE patients. NEK7 seems to be involved in the recruitment of centrosomal pericentriolar proteins that are necessary for centriole duplication and spindle pole formation during mitosis.^80^ Chromosomal lagging, micronuclei formation, cytokinesis failure and tetraploidy/aneuploidy were observed in NEK7 deficient mouse embryonic fibroblasts.^81^ Therefore, deregulation of NEK7 activity in cell division processes might contribute to aberrant mitosis and induction of the relapse status.^82^ Furthermore, we found two phosphosites, S372 and S389 in the ribosomal S6 kinase RPS6KA1 isoform 2 (Uniprot identifier: Q15418-2) upregulated in REL_FREE patients. It has been reported that phosphorylation at S372 and S389, possibly by PRKC activators, switched on the RPS6KA1 N-terminal kinase domain required for phosphorylation of its substrates.^83^ The role of ribosomal S6 kinases in cancer is not well understood and seems to vary by cancer type. While some isoforms of ribosomal S6 kinases can promote cell motility and invasion processes by altering transcription and integrin activity, others depress them by interacting with the actin cytoskeleton.^84^ The fact that the phosphosites of RPS6KA1 might be phosphorylated by NEK7^82^, and not by MTOR, might explain the benefits of higher phosphorylation of membrane, actin filament and cell-cell adherens junction proteins in the REL_FREE group. Our findings encourage functional studies to understand the role of NEK7, PRKC and RPS6KA1 kinases in the cytoskeleton of AML cells.

In conclusion, the proteomics profiles of patient-derived AML cells collected at the first time of diagnosis differ between patients that are long-term relapse free survivors and patients that relapse. The relapse-associated group displayed decreased expression of V-ATPases and increased activity of CDKs and CSK2. Additionally, our results suggest a subset of transcriptional and metabolic regulators that could be considered as predictors of prognosis and therapeutic targets in AML.

## Acknowledgments

Hilde Kristin Garberg, Olav Mjaavatten, Marie Hagen, Kristin Rye Paulsen, Atle Brendehaug, Sigrid Erdal, Laura Minsaas, Hans Petter Brodahl and Nina Lied Larsen for excellent technical assistance. The Genomics Core Facility (GCF) at the University of Bergen, which is part of the NorSeq consortium, provided support in ChIP-Seq bioinformatics analysis; GCF is supported in part by major grants from the Research Council of Norway (grant no. 245979/F50) and Bergen Research Foundation (BFS). Kreftforeningen (the Norwegian Cancer Society, grant no. 100933) for financial support. Work at The Novo Nordisk Foundation Center for Protein Research (CPR) is funded in part by a generous donation from the Novo Nordisk Foundation (Grant number NNF14CC0001).

## Authorship

Contribution: EA performed research, analyzed proteomics data and wrote the paper. MHV contributed to experimental design, performed research, analyzed phosphoproteomics data and wrote the paper. FB and FS contributed to scientific discussions, experimental design and paper writing. SSB performed cell proliferation assays. ØB provided the AML patient samples and contributed to scientific discussions, experimental design and paper writing. TS performed ChIP-seq analyses. RH performed mutational studies of the AML patients. SOD contributed to scientific discussions and paper writing. EMC and MV contributed to scientific discussions. TSB contributed to the data analysis, scientific discussions and paper writing. JVO contributed to the data analysis, scientific discussions and input on paper writing.

